# Toxicity of House Cricket (*Acheta domesticus*) in Mice

**DOI:** 10.1101/2022.06.23.497304

**Authors:** Matano Yasuki, Sakagami Kiyo, Nojiri Yuuto, Nomura Kenta, Masuda Akira, Moriike Yuuki, Yamamoto Akane, Nagai Nobuo, Ogura Atsushi

**Author notes:** Corresponding author: Nagai Nobuo, Affiliation address: Department of Animal Bioscience, Nagahama Institute of Bio-Science and Technology, Tamura 1266, Nagahama, Shiga 526-0829, JAPAN.

## Abstract

There is an urgent need to address the shortage of animal protein due to food shortages caused by the global population growth. Crickets contain an abundance of proteins in their exoskeleton and muscles and have attracted attention as a new protein source; however, their safety as a food source has not been confirmed. We evaluated the toxicity of the House cricket (*Acheta domesticus*), on cells and mammals. In genotoxicity *in vitro*, cricket powder was added to Chinese hamster lung CHL-IU cells at concentrations of 5,000 µg/mL, and the rate of chromosomal aberrations was assessed. In genotoxicity *in vivo*, mice were orally administered up to 2,000 mg/kg of cricket powder for 2 days. In both tests, cricket powder did not show any toxic effect. A repeated oral toxicity study was performed administering up to 3,000 mg/kg of cricket powder or control (saline) for 14 or 90 consecutive days and measuring body weight changes, blood biochemistry, blood properties, and organ weights. In each time course, there were no differences in there parameters between the control and cricket powder treated groups. These results suggest that House crickets (≤3,000 mg/kg) are not toxic to cells and organisms.

## Introduction

The world’s population is predicted to grow to 9 billion by 2030 and solutions to the problems of food shortage caused by this population growth are required urgently. There are concerns that the demand for animal protein will increase rapidly and lead to deforestation due to increased production and overgrazing of livestock, such as cattle and pigs, and exacerbation of environmental problems caused by greenhouse gases, such as methane and carbon dioxide, emitted from livestock (Oonincx *et al*., 2010). One possible solution to address these food problems is an insect diet, which is high in protein, rich in minerals, and has high nutritional value. Insects are already used as food in many parts of the world, including Asia, North and South America, and Africa, and it is estimated that 1,400 species of insects are used as food worldwide. Furthermore, insects are considered to have a lower environmental impact than livestock, such as cattle and pigs, because they can be raised on smaller areas of land, require less food and water, and emit less methane and carbon dioxide gas (Nowakowski *et al*., 2021). It has been reported that insects contain many bioactive substances beneficial to humans, such as vitamin C, polyphenols, and glycosaminoglycans, which have antioxidant and anti-inflammatory properties (Di Mattia *et al*., 2019; Zielińska *et al*., 2017; Ahn *et al*., 2020).

The House cricket (*Acheta domesticus*) is an insect belonging to the family of crickets in the order Grasshopperae. It is commercially bred for use as food for amphibians, birds, and reptiles. Crickets are composed of protein and fat (e.g., in the exoskeleton and muscle) and are considered to be a high-quality food source of animal protein (Huis, 2013). However, the toxicity and safety of insects as food remains unclear (Klunder *et al*., 2012). Therefore, we evaluated the toxicity of dried house cricket powder in cells and mammals. Genotoxicity was evaluated by an *in vitro* chromosome aberration test using Chinese hamster lung-derived CHL-IU cells and by an *in vivo* micronucleus test using mice. In addition, a repeated dose 14 and 90 consecutive days oral toxicity study was performed in mice.

EFSA Panel on Nutrition, Novel Food and Food Allergens (NDA) concludes that the house crickets (Acheta domesticus) is safe under the three formulations: frozen, dried, and ground (Turck, *et al*., 2021a). Following this conclusion, the European Commission authorised the marketing of house crickets (Acheta domesticus) as novel food in the EU as of 11 February 2022. The Panel, however, only discusses toxicological information based on various aspects, but does not test for genotoxicity and just referred that toxicological test on the G. bimaculatus. However, they notes that G. bimaculatus belongs to the same family as A. domesticus (Gryllidae), but these are different species and also the rearing conditions of the insects used in the study are not known. Therefore, the addition of toxicological tests in house cricket in this paper is very meaningful in supporting the marketing of house cricket.

## Materials and methods

### Preparation of cricket powder

House cricket powder was provided by Ecologgie Inc. (Tokyo, Japan) and used for all safety studies. Two-day fasted House crickets were euthanized in ice and washed thoroughly in running water. Next, 100-200 g of dried crickets was boiled in water at 100°C for 15 min, dried at 70°C for 12 hr to a moisture content of 5%-7%, and crushed for 30 s five times to produce dried cricket powder. The cricket powder was also sterilized (KOGA ISOTOPE Inc. Shiga, Japan) using 30 Gy γ-rays and used for experiments. For the animal experiments, cricket powder was suspended in 0.9% saline and administered orally at various concentrations.

### Animals

All animal experimental procedures were approved by the Committee on Animal Care and Use of the Nagahama Institute of Bio-Science and Technology (Permit Number: 099). The animal studies were performed in accordance with the institutional and national guidelines and regulations, and with the ARRIVE (Animal Research: Reporting of In Vivo Experiments) guidelines (https://www.nc3rs.org.uk/arrive-guidelines). An *in vivo* micronucleus test and 14 and 90 consecutive days oral toxicity study with a repeated dose were performed according to the Organization for Economic Cooperation and Development (OECD) guidelines No. 474 (https://doi.org/10.1787/9789264264762-en) and 408 (https://doi.org/10.1787/9789264070707-en), respectively. JcI:ICR female and male mice (CLEA Japan, Tokyo, Japan) aged 5 weeks were used. These animals were maintained at approximately 23°C, 50% humidity, 18.6 m^3^/min on charge and 17.1 m^3^/min on return in air ventilation, and a 12-hr light/dark cycle. Mice were fed a rodent diet *ad libitum* (CE-2; CLEA Japan, Tokyo, Japan). The number of animals used in each experiment is described in each figure legend.

### Genotoxicity analysis

#### Mammalian chromosomal aberration test

A mammalian chromosomal aberration test for cricket power was performed according to OECD guideline 473 for testing chemicals (https://doi.org/10.1787/9789264264649-en). CHL-IU cells were plated in E-MEM (Fujifilm, 051-07615, Tokyo, Japan) containing heat-inactivated 10% fetal bovine serum at a density of 4 × 10^4^ / 6 cm dish and incubated for 48 hr. Positive control cells were treated with a combination of metabolic activator S9-mix (Ieda boeki, Tokyo, Japan) and 200 ng/mL benzo[a]pyrene (MilliporeSigma, B1790, Saint Louis, USA). Negative control cells were treated without metabolic activator and/or benzo[a]pyrene. The toxicity of house cricket powder was examined under three conditions as follows: 5,000 µg of cricket powder/mL without S9-mix for 6 hr; 5,000 µg of cricket powder/mL with S9-mix for 22 hr; and 5,000 µg of cricket powder/mL without S9-mix for 22 hr.

For the 6-hr treatment and 16-hr recovery incubation, cells were rinsed and the media were replaced with fresh media after the 6 hr treatment, followed by culture for an additional 16 hr. Colcemid (Fijifilm 47253, Tokyo, Japan) at final 100 ng/µL was added for the last 2 hr and then cells were harvested with 0.25% trypsin/ 1 mM EDTA (Nacalai Tesque, 32777, Kyoto, Japan). Single cells were treated with 75 mM KCl hypotonic solution for 20 min at 37°C and fixed with Carnoy’s solution (3:1=methanol: acetic acid). Giemsa staining was performed in specimens after fixation with methanol for 3 min. Staining was performed using 2% Giemsa solution (Merck, 1.109204, Kenilworth, USA)/Giemsa buffer (pH6.8) for 5 min at room temperature followed by washing in running water for 2 min. Air-dried samples were mounted in Multi Mount 480 (Matsunami Glass, Osaka, Japan) and more than 100 cells were observed under a microscope (Olympus CKX53, Tokyo, Japan).

#### *In vivo* micronucleus test

ICR mice were acclimated for 1 week in the breeding room and then checked for health and weight. Mice were divided into five groups (5-7 animals/sex/group) and treated with 500, 1,000, or 2,000 mg/kg of cricket powder, 0.9% saline (control), or cyclophosphamide monohydrate (CPA; Sigma Chemical Company, Saint Louis, USA) as a positive control (Han *et al*., 2014). Cricket powder and control groups received two oral doses at 24-hr intervals, while the CPA group received a single intraperitoneal dose of 70 mg/kg CPA. Under deep anesthesia by intraperitoneal administration of 200 mg/kg sodium secobarbital solution (Nichi-Iko Pharmaceutical Company, Toyama, Japan) 24 hr after the last dose, both femur were removed. Then bone marrow fluid was extracted from the femur and centrifuged at 1,200 rpm for 5 min to isolate bone marrow cells, and their smears were prepared on glass slides (Schmid, 1975). After drying at room temperature, smears were fixed with methanol and stained for nucleic acids using acridine orange solution (Fujifilm, Osaka, Japan). The percentage of micronucleated (MN) polychromatic erythrocytes (PCEs) in 2,000 juvenile erythrocytes per mouse, including PCEs and normochromatic erythrocytes (NCEs), was calculated. The ratio of PCEs per 500 erythrocytes [PCEs/(PCEs+NCEs)] was measured to assess genotoxicity and cytotoxicity.

#### Oral toxicity studies for 14- and 90-day repeated dosing

In the repeated dose 14-day oral toxicity study, ICR mice were acclimated for 1 week in the breeding room, weighed, and divided into four groups of 4-9 animals/sex/group to receive treatment of 300, 1,000, or 3,000 mg/kg cricket powder or 0.9% saline (control). Mice were orally administered cricket powder or saline for 14 days, weighed, checked daily for abnormalities in condition and death. Mice were fasted for no more than 16 hr after last administration on the 14th day. Under anesthesia by intraperitoneal administration of 200 mg/kg sodium secobarbital solution, blood characterization, blood biochemical tests, and organ weights were assessed. Blood biochemistry tests were performed by serum prepared from blood collected from a posterior vena cava in each animal, coagulated at 37°C for at least 2 hr, and centrifuged at 3,300 rpm (4°C) for 15 min to extract the serum. The serum was analyzed using Fuji Drychem 7000 (Fujifilm, Osaka, Japan) to measure levels of aspartate aminotransferase (AST), alanine aminotransferase (ALT), alkaline phosphatase (ALP), and total bilirubin (T-BIL), blood urea nitrogen (BUN), creatinine (CRE), uric acid (UA), glucose (GLU), total cholesterol (CHO), triglycerides (TG), and total protein (TP), albumin (ALB), lactate dehydrogenase (LDH), creatine phosphokinase (CPK), calcium (Ca), inorganic phosphorus (IP), magnesium (Mg), sodium (Na), potassium (K), and chloride (Cl).

Blood characterization was performed by collecting approximately 100 μL of blood from each animal via vena cava using EDTA as an anticoagulant. The numbers of white blood cell (WBC), red blood cell (RBC), and total blood cell counts together with hemoglobin concentration, hematocrit ratio, mean corpuscular volume, mean corpuscular hemoglobin (MCH), RBC distribution width, platelets, and leukocytes (lymphocyte, monocyte, granulocyte, eosinocyte count and ratio) were performed using an LC-662 blood cell counter (FUKUDA DENSHI, Shiga, Japan).

Mice were euthanized and the brain, pituitary gland, submandibular gland, thymus, heart, lung, liver, spleen, pancreas, adrenal gland (right and left), kidney (right and left), ovary (right and left), testis (right and left), and seminal vesicle/coagulating gland were collected and weighed using an electronic balance. Each organ weight was converted to percentage of total body weight after fasting. These results were used to determine the dose for the 90-day oral toxicity study.

In the repeated dose 90-day oral toxicity study, mice were orally administered cricket powder or saline then weighed, checked daily for abnormalities in condition and death for 90 days. The mice were then fasted for 16 hr after the 90th day dose, deeply anesthetized by intraperitoneally administration of 200 mg/kg of sodium secobarbital solution. Blood measurements were made as per the 14-day oral toxicity study. Blood biochemistry was assessed for ALT, BUN, CRE, CHO, TG, ALB, CPK, Ca, Mg, and Cl. In addition to the organs in the 14-day oral toxicity study, uterus, visceral fat (right and left), subcutaneous fat, gastrocnemius muscle (right and left), and soleus muscle (right and left) were measured and expressed as percentage of total body weight after fasting.

### Statistical analysis

Statistical analyses were performed using Fisher’s exact test for *in vitro* chromosomal aberration test, and one-way analysis of variance followed by Dunnet test for the analysis of the *in vivo* micronucleus test and oral toxicity study. *P*-values <0.05 were considered significant.

## Results

### *In vitro* chromosomal aberration test

The effect of cricket powder at 5,000 µg/mL on CHL-IU cells was analyzed on in vitro chromosomal aberration test. Negative control without Benzo[a]pyrene and/or S9-mix showed 3∼4% numerical aberration. Positive control with Benzo[a]pyrene at 200 ng/mL resulted in 15.7% of numerical aberration which is polyploid, endoreduplicated or hyperdiploid cells, which showed a significant increase (*P* < 0.01) compared to the negative control. In contrast, there was not significant difference in numerical aberration between cells treated with 5,000 µg/mL cricket powder compared with the negative control (Table 1). These results indicate that 5,000 µg/mL house cricket powder has no cytotoxicity on CHL-IU cells.

**Table 1.** Mammalian chromosomal aberration test.

| Substances | Dose | Treatment-recovery<br>time (hours) | S9 mix | Frequency of cells with<br>numerical aberration<br>(%) | Number of<br>metaphase |
| --- | --- | --- | --- | --- | --- |
| DMSO | n/a | 6-16 | – | 3.3 | 122 |
| DMSO | n/a | 6-16 | + | 3.4 | 116 |
| Benzo[a]pyrene | 200 ng/mL | 6-16 | – | 3.1 | 126 |
| Benzo[a]pyrene | 200 ng/mL | 6-16 | + | 12.7** | 126 |
| cricket powder | 5000 µg/mL | 6-16 | – | 5.4 | 128 |
| cricket powder | 5000 µg/mL | 22-0 | – | 8.6 | 116 |
| cricket powder | 5000 µg/mL | 22-0 | + | 4.5 | 111 |

\*\* $P < 0.01$ vs DMSO without S9 mix.

### *In vivo* micronucleus analysis

There was no difference in the ratio of MNPCEs per 2,000 PCEs and PCEs per 500 erythrocytes in the 500, 1,000, and 2,000 mg/kg cricket powder treatment groups compared with the control group for both sexes. On the other hand, there was a significant increase (*P* < 0.01) in the percentage of MNPCEs per 2,000 PCEs in the CPA group compared with the control group, but no difference in the percentage of PCEs per 500 erythrocytes (Table 2). These results suggest that ≤2,000 mg/kg House cricket powder is not genotoxic or cytotoxic in mice.

**Table 2.**
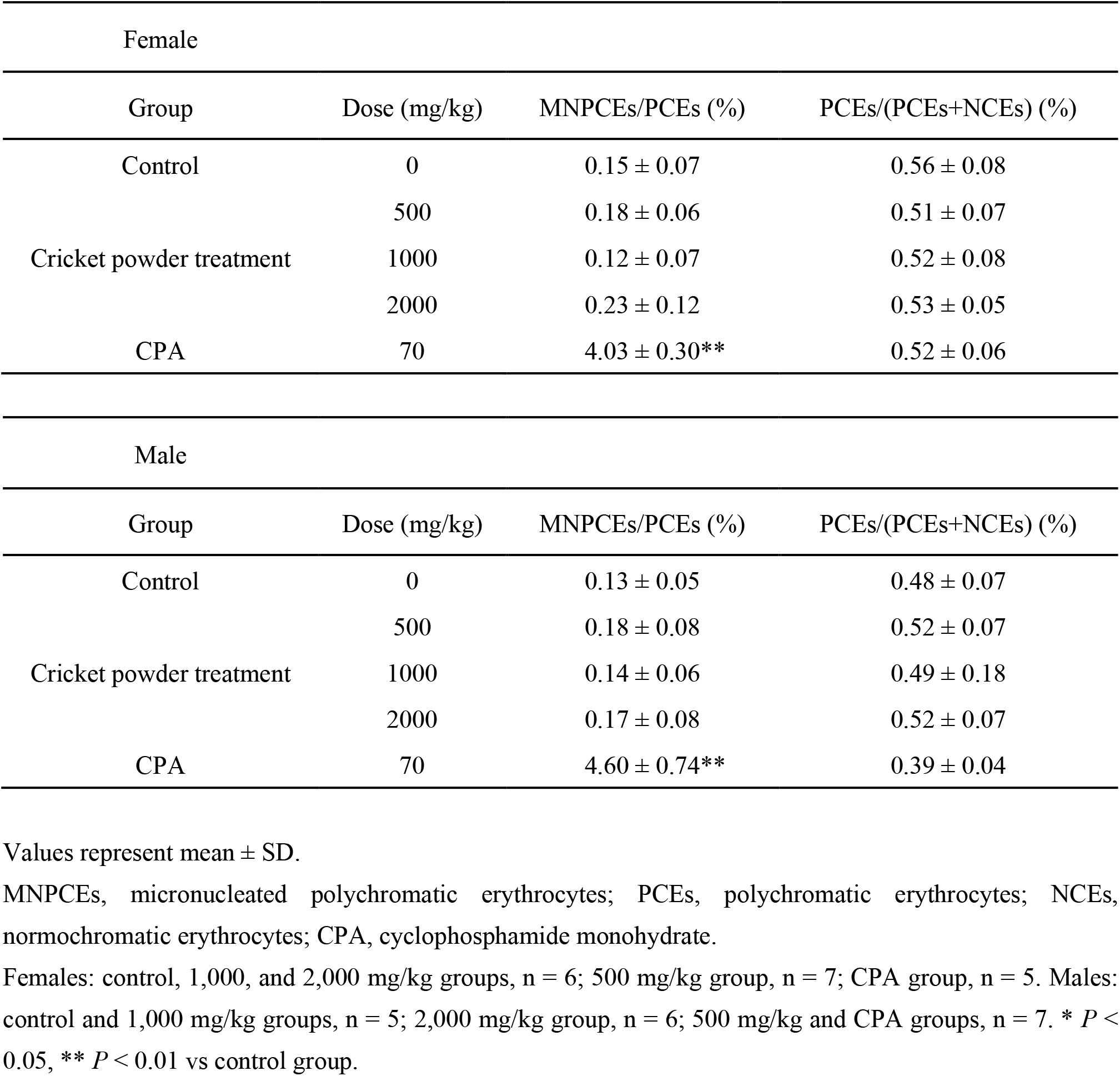
*In vivo* micronucleus test.

### Repeated dose for 14- and 90-day oral toxicity studies

In the 14-day oral toxicity study, there were no differences in body weight changes or blood biochemistry tests during the 14-day treatment period in the 300, 1,000, and 3,000 mg/kg cricket powder groups compared with the control group for both males and females (Fig. 1 and Table 3. Blood biochemistry tests revealed a significant increase (*P* < 0.05) in MCH in the female 1,000 mg/kg group compared with the control group, but no difference in males in the cricket powder group compared with the control group (Table 4). In addition, there was no difference in the ratio of organ to body weight after fasting in male and female mice treated with cricket powder compared with the control group (Table 5).

**Table 3.** Serum biochemical values of mice treated with House cricket powder for 14 days.

| Female |  |  |  |  |
| --- | --- | --- | --- | --- |
|  | Group |  |  |  |
|  | Control | 300 mg/kg | 1000 mg/kg | 3000 mg/kg |
| AST<sup>1</sup> (IU/L) | 66 ± 31 | 68 ± 9 | 76 ± 25 | 60 ± 33 |
| ALT<sup>2</sup> (IU/L) | 23 ± 4 | 24 ± 6 | 24 ± 6 | 23 ± 8 |
| ALP<sup>3</sup> (IU/L) | 76 ± 18 | 100 ± 28 | 106 ± 32 | 81 ± 3 |
| T-BIL<sup>4</sup> (mg/dL) | 0.6 ± 0.11 | 0.7 ± 0.10 | 0.5 ± 0.04 | 0.5 ± 0.05 |
| BUN<sup>5</sup> (mg/dL) | 26.6 ± 3.91 | 24.4 ± 6.72 | 24.2 ± 6.31 | 23.2 ± 5.01 |
| CRE<sup>6</sup> (mg/dL) | 0.13 ± 0.04 | 0.14 ± 0.04 | 0.11 ± 0.03 | 0.12 ± 0.03 |
| UA<sup>7</sup> (mg/dL) | 1.4 ± 0.26 | 2.1 ± 1.17 | 1.4 ± 0.22 | 1.8 ± 0.75 |
| GLU<sup>8</sup> (mg/dL) | 149 ± 49 | 172 ± 83 | 131 ± 39 | 155 ± 38 |
| CHO<sup>9</sup> (mg/dL) | 73 ± 7 | 81 ± 15 | 77 ± 15 | 81 ± 18 |
| TG<sup>10</sup> (mg/dL) | 98 ± 32 | 128 ± 48 | 101 ± 24 | 46 ± 21 |
| TP<sup>11</sup> (g/dL) | 5.3 ± 0.11 | 5.5 ± 0.10 | 5.0 ± 0.23 | 5.3 ± 0.31 |
| ALB<sup>12</sup> (g/dL) | 2.4 ± 0.11 | 2.6 ± 0.12 | 2.5 ± 0.17 | 2.6 ± 0.28 |
| LDH<sup>13</sup> (IU/L) | 585 ± 241 | 814 ± 163 | 761 ± 172 | 624 ± 200 |
| CPK<sup>14</sup> (U/L) | 86 ± 25 | 136 ± 49 | 175 ± 50 | 120 ± 63 |
| Ca<sup>15</sup> (mg/dL) | 10.1 ± 0.42 | 10.3 ± 0.64 | 9.6 ± 0.48 | 10.3 ± 1.28 |
| IP<sup>16</sup> (mg/dL) | 13.1 ± 1.77 | 14.2 ± 1.49 | 13.5 ± 0.83 | 14.4 ± 0.76 |
| Mg<sup>17</sup> (mg/dL) | 3.5 ± 0.58 | 4.0 ± 0.71 | 3.6 ± 0.29 | 3.7 ± 0.68 |
| Na<sup>18</sup> (mmol/L) | 150 ± 1 | 150 ± 2 | 151 ± 1 | 152 ± 1 |
| K<sup>19</sup> (mmol/L) | 5.8 ± 1.30 | 7.7 ± 2.36 | 5.4 ± 0.89 | 6.4 ± 1.51 |
| Cl<sup>20</sup> (mmol/L) | 113 ± 4 | 115 ± 3 | 114 ± 4 | 117 ± 3 |

Male
|  | Group |  |  |  |
| --- | --- | --- | --- | --- |
|  | Control | 300 mg/kg | 1000 mg/kg | 3000 mg/kg |
| AST<sup>1</sup> (IU/L) | 66 ± 17 | 47 ± 22 | 54 ± 12 | 40 ± 6 |
| ALT<sup>2</sup> (IU/L) | 29 ± 10 | 19 ± 4 | 22 ± 4 | 22 ± 4 |
| ALP<sup>3</sup> (IU/L) | 73 ± 18 | 68 ± 23 | 69 ± 16 | 65 ± 21 |
| T-BIL<sup>4</sup> (mg/dL) | 0.6 ± 0.12 | 0.6 ± 0.15 | 0.5 ± 0.08 | 0.4 ± 0.10 |
| BUN<sup>5</sup> (mg/dL) | 30.2 ± 7.68 | 25.8 ± 3.72 | 24.1 ± 3.98 | 30.6 ± 7.88 |
| CRE<sup>6</sup> (mg/dL) | 0.16 ± 0.07 | 0.11 ± 0.02 | 0.10 ± 0.01 | 0.14 ± 0.04 |
| UA<sup>7</sup> (mg/dL) | 2.0 ± 0.79 | 1.5 ± 0.28 | 1.9 ± 0.77 | 1.3 ± 0.29 |
| GLU<sup>8</sup> (mg/dL) | 156 ± 46 | 119 ± 28 | 184 ± 41 | 162 ± 25 |
| CHO<sup>9</sup> (mg/dL) | 111 ± 16 | 124 ± 23 | 115 ± 4 | 101 ± 19 |
| TG<sup>10</sup> (mg/dL) | 69 ± 20 | 77 ± 23 | 100 ± 24 | 62 ± 34 |
| TP<sup>11</sup> (g/dL) | 5.4 ± 0.16 | 5.3 ± 0.64 | 5.2 ± 0.11 | 4.8 ± 0.19 |
| ALB<sup>12</sup> (g/dL) | 2.5 ± 0.10 | 2.5 ± 0.44 | 2.5 ± 0.11 | 2.2 ± 0.12 |
| LDH<sup>13</sup> (IU/L) | 729 ± 210 | 586 ± 263 | 517 ± 116 | 405 ± 111 |
| CPK<sup>14</sup> (U/L) | 194 ± 60 | 159 ± 111 | 107 ± 23 | 122 ± 31 |
| Ca<sup>15</sup> (mg/dL) | 10.2 ± 0.39 | 10.1 ± 0.63 | 10.3 ± 0.72 | 9.7 ± 0.32 |
| IP<sup>16</sup> (mg/dL) | 14.1 ± 1.32 | 14.0 ± 1.20 | 14.1 ± 0.25 | 12.3 ± 2.27 |
| Mg<sup>17</sup> (mg/dL) | 4.0 ± 0.81 | 4.2 ± 1.29 | 3.7 ± 0.27 | 3.6 ± 0.26 |
| Na<sup>18</sup> (mmol/L) | 151 ± 0.4 | 151 ± 1 | 150 ± 2 | 151 ± 1 |
| K<sup>19</sup> (mmol/L) | 7.2 ± 1.69 | 5.5 ± 0.89 | 6.5 ± 1.41 | 5.0 ± 0.79 |
| Cl<sup>20</sup> (mmol/L) | 113 ± 1 | 112 ± 2 | 113 ± 2 | 111 ± 2 |

Values represent mean ± SD.
1: Aspartate aminotransferase, 2: Alanine aminotransferase, 3: alkaline phosphatase, 4: total bilirubin, 5: blood urea nitrogen, 6: creatinine, 7: uric acid, 8: glucose, 9: total cholesterol, 10: triglycerides, 11: total protein, 12: albumin, 13: lactate dehydrogenase, 14: creatine phosphokinase, 15: calcium, 16: inorganic phosphorus, 17: magnesium, 18: sodium, 19: potassium, and 20: chloride.
Females: control group, n = 4; 300, 1,000, and 3,000 mg/kg groups, n = 5. Males: control and 1,000 mg/kg groups, n = 4; 300 and 3,000 mg/kg groups, n = 5.

**Table 4.** Hematological values of mice treated with House cricket powder for 14 days.

| Female |  |  |  |  |
| --- | --- | --- | --- | --- |
|  | Group |  |  |  |
|  | Control | 300 mg/kg | 1000 mg/kg | 3000 mg/kg |
| WBC<sup>1</sup> (10<sup>3</sup>/μL) | 7.05 ± 2.25 | 9.65 ± 2.07 | 7.05 ± 3.37 | 7.65 ± 3.50 |
| RBC<sup>2</sup> (10<sup>6</sup>/μL) | 6.77 ± 1.83 | 7.88 ± 1.23 | 5.96 ± 0.78 | 7.49 ± 1.10 |
| Hemoglobin (g/dL) | 10.10 ± 2.54 | 12.83 ± 1.77 | 10.25 ± 1.47 | 12.40 ± 2.03 |
| Hematocrit (%) | 32.98 ± 9.67 | 40.15 ± 6.09 | 30.78 ± 5.96 | 37.73 ± 6.53 |
| MCV<sup>3</sup> (μm<sup>3</sup>) | 48.43 ± 1.27 | 50.98 ± 0.39 | 51.23 ± 3.58 | 50.20 ± 1.71 |
| MCH<sup>4</sup> (pg) | 15.03 ± 1.11 | 16.30 ± 0.37 | 17.20 ± 0.78* | 16.55 ± 0.52 |
| MCHC<sup>5</sup> (g/dL) | 31.00 ± 2.39 | 31.95 ± 0.59 | 33.75 ± 2.41 | 33.00 ± 0.61 |
| RDW<sup>6</sup> (%) | 14.05 ± 0.51 | 13.53 ± 0.47 | 15.05 ± 2.36 | 14.00 ± 0.46 |
| Platelets (10<sup>3</sup>/μL) | 420.8 ± 340.5 | 146.0 ± 79.6 | 304.0 ± 35.4 | 113.5 ± 83.7 |
| Lymphocyte (10<sup>3</sup>/μL) | 3.18 ± 1.33 | 4.83 ± 0.64 | 3.65 ± 1.47 | 4.00 ± 2.33 |
| Monocyte (10<sup>3</sup>/μL) | 0.23 ± 0.08 | 0.30 ± 0.07 | 0.20 ± 0.07 | 0.23 ± 0.11 |
| Granulocyte (10<sup>3</sup>/μL) | 3.65 ± 1.87 | 4.53 ± 1.63 | 3.20 ± 1.87 | 3.43 ± 1.16 |
| Eosinocyte (10<sup>3</sup>/μL) | 0.90 ± 0.63 | 1.40 ± 0.79 | 0.98 ± 0.63 | 1.33 ± 0.36 |

| Male |  |  |  |  |
| --- | --- | --- | --- | --- |
|  | Group |  |  |  |
|  | Control | 300 mg/kg | 1000 mg/kg | 3000 mg/kg |
| WBC<sup>1</sup> (10<sup>3</sup>/μL) | 5.85 ± 2.36 | 3.94 ± 1.92 | 6.15 ± 2.25 | 9.24 ± 5.12 |
| RBC<sup>2</sup> (10<sup>6</sup>/μL) | 6.07 ± 2.01 | 6.73 ± 1.30 | 7.11 ± 0.28 | 6.33 ± 2.17 |
| Hemoglobin (g/dL) | 9.88 ± 3.36 | 10.88 ± 2.03 | 11.40 ± 0.48 | 10.31 ± 3.55 |
| Hematocrit (%) | 30.86 ± 10.90 | 33.32 ± 7.64 | 35.43 ± 1.38 | 33.05 ± 11.14 |
| MCV<sup>3</sup> (μm<sup>3</sup>) | 50.68 ± 1.63 | 49.14 ± 2.89 | 49.85 ± 0.32 | 52.38 ± 1.60 |
| MCH<sup>4</sup> (pg) | 16.21 ± 0.28 | 16.16 ± 0.48 | 16.05 ± 0.38 | 16.26 ± 0.59 |
| MCHC<sup>5</sup> (g/dL) | 32.01 ± 1.14 | 33.04 ± 1.65 | 32.23 ± 0.75 | 31.03 ± 0.57 |
| RDW<sup>6</sup> (%) | 14.26 ± 0.55 | 14.36 ± 1.80 | 13.60 ± 0.39 | 13.58 ± 0.61 |
| Platelets (10<sup>3</sup>/μL) | 324.6 ± 171.9 | 256.4 ± 181.5 | 310.5 ± 253.1 | 172.0 ± 210.7 |
| Lymphocyte (10<sup>3</sup>/μL) | 2.90 ± 1.50 | 1.58 ± 1.15 | 3.05 ± 1.02 | 4.16 ± 2.03 |
| Monocyte (10<sup>3</sup>/μL) | 0.19 ± 0.08 | 0.12 ± 0.04 | 0.15 ± 0.05 | 0.24 ± 0.12 |
| Granulocyte (10<sup>3</sup>/μL) | 2.76 ± 1.19 | 2.24 ± 0.80 | 2.95 ± 1.24 | 4.84 ± 3.60 |
| Eosinocyte (10<sup>3</sup>/μL) | 1.05 ± 0.80 | 0.90 ± 0.34 | 0.98 ± 0.49 | 1.96 ± 1.68 |

Values represent mean ± SD.
1: White blood cell, 2: Red blood cell, 3: Mean corpuscular volume, 4: Mean corpuscular hemoglobin, 5: Mean corpuscular hemoglobin concentration, and 6: RBC distribution width.
Females: n = 4 for all groups. Males: control group, n = 8; 300 and 1,000 mg/kg groups, n = 4; 3,000 mg/kg group, n = 9. \* $P < 0.05$ vs. control group.

**Table 5.** Relative organ weight of mice treated with House cricket powder for 14 days.

| Female |  |  |  |  |
| --- | --- | --- | --- | --- |
| OW/BW (%) | Group |  |  |  |
|  | Control | 300 mg/kg | 1000 mg/kg | 3000 mg/kg |
| Brain | 1.5408 ± 0.1810 | 1.6776 ± 0.1124 | 1.5705 ± 0.1279 | 1.7254 ± 0.1105 |
| Pituitary gland | 0.0079 ± 0.0023 | 0.0073 ± 0.0015 | 0.0087 ± 0.0060 | 0.0118 ± 0.0051 |
| Submandibular gland | 0.6025 ± 0.1620 | 0.5426 ± 0.0936 | 0.6426 ± 0.0988 | 0.6511 ± 0.0831 |
| Thymus | 0.2887 ± 0.0499 | 0.3076 ± 0.0730 | 0.2984 ± 0.0464 | 0.2982 ± 0.1014 |
| Heart | 0.4688 ± 0.0315 | 0.5179 ± 0.0419 | 0.5022 ± 0.0497 | 0.5229 ± 0.0334 |
| Lung | 0.6220 ± 0.0195 | 0.6373 ± 0.0996 | 0.6548 ± 0.1077 | 0.6353 ± 0.0701 |
| Liver | 5.0887 ± 0.3949 | 4.9985 ± 0.5928 | 5.5440 ± 0.9749 | 4.8743 ± 0.1564 |
| Spleen | 0.3862 ± 0.1518 | 0.4001 ± 0.0849 | 0.3359 ± 0.0682 | 0.3257 ± 0.0481 |
| Pancreas | 0.9261 ± 0.3614 | 0.9658 ± 0.2838 | 1.0538 ± 0.6274 | 0.8304 ± 0.1981 |
| Adrenal gland (R) | 0.0161 ± 0.0055 | 0.0202 ± 0.0133 | 0.0152 ± 0.0047 | 0.0291 ± 0.0218 |
| Kidney (R) | 0.6978 ± 0.0461 | 0.7007 ± 0.0826 | 0.7403 ± 0.0685 | 0.7397 ± 0.0648 |
| Adrenal gland (L) | 0.0197 ± 0.0084 | 0.0190 ± 0.0054 | 0.0199 ± 0.0045 | 0.0256 ± 0.0052 |
| Kidney (L) | 0.6631 ± 0.0356 | 0.6592 ± 0.1161 | 0.6972 ± 0.0647 | 0.7382 ± 0.0592 |
| Ovary (R) | 0.0182 ± 0.0087 | 0.0157 ± 0.0018 | 0.0251 ± 0.0094 | 0.0285 ± 0.0101 |
| Ovary (L) | 0.0219 ± 0.0074 | 0.0247 ± 0.0125 | 0.0240 ± 0.0063 | 0.0290 ± 0.0113 |

Male
| OW/BW (%) | Group |  |  |  |
| --- | --- | --- | --- | --- |
|  | Control | 300 mg/kg | 1000 mg/kg | 3000 mg/kg |
| Brain | 1.3045 ± 0.0905 | 1.3570 ± 0.2318 | 1.2107 ± 0.0751 | 1.3151 ± 0.1686 |
| Pituitary gland | 0.0058 ± 0.0020 | 0.0092 ± 0.0051 | 0.0060 ± 0.0029 | 0.0063 ± 0.0019 |
| Submandibular gland | 0.6856 ± 0.0703 | 0.7845 ± 0.2069 | 0.6229 ± 0.0958 | 0.6671 ± 0.0953 |
| Thymus | 0.1517 ± 0.0398 | 0.1302 ± 0.0452 | 0.1608 ± 0.0306 | 0.1317 ± 0.0484 |
| Heart | 0.4863 ± 0.0468 | 0.4786 ± 0.0717 | 0.4877 ± 0.0355 | 0.4936 ± 0.0628 |
| Lung | 0.6517 ± 0.0631 | 0.5867 ± 0.0724 | 0.6279 ± 0.1379 | 0.5776 ± 0.0547 |
| Liver | 4.7717 ± 0.3678 | 4.5986 ± 0.7985 | 4.8139 ± 0.2388 | 4.7760 ± 0.4950 |
| Spleen | 0.2756 ± 0.0921 | 0.2320 ± 0.0358 | 0.2433 ± 0.0463 | 0.2770 ± 0.0594 |
| Pancreas | 0.6780 ± 0.2863 | 0.7191 ± 0.3076 | 0.9481 ± 0.2257 | 0.6499 ± 0.1727 |
| Adrenal gland (R) | 0.0078 ± 0.0032 | 0.0100 ± 0.0051 | 0.0087 ± 0.0076 | 0.0100 ± 0.0046 |
| Kidney (R) | 0.8610 ± 0.0523 | 0.8884 ± 0.1110 | 0.8878 ± 0.0844 | 0.9230 ± 0.0603 |
| Adrenal gland (L) | 0.0109 ± 0.0035 | 0.0104 ± 0.0066 | 0.0062 ± 0.0034 | 0.0101 ± 0.0037 |
| Kidney (L) | 0.8228 ± 0.0724 | 0.8228 ± 0.0716 | 0.8421 ± 0.0540 | 0.8888 ± 0.0693 |
| Testis (R) | 0.4301 ± 0.0369 | 0.4551 ± 0.0466 | 0.3894 ± 0.0161 | 0.4015 ± 0.0500 |
| Testis (L) | 0.3926 ± 0.0498 | 0.4496 ± 0.0402 | 0.3767 ± 0.0087 | 0.3795 ± 0.0564 |
| Seminal vesicle/Coagulating gland | 0.7160 ± 0.1280 | 0.7586 ± 0.1687 | 0.7665 ± 0.0398 | 0.7614 ± 0.1505 |

Values represent mean ± SD.
OW, organ weight; BW, body weight.
Females: all groups, n = 5. Males: control group, n = 8; 300 mg/kg group, n = 5; 1,000 mg/kg group, n = 4; 3,000 mg/kg group, n = 9.

**Figure 1.**
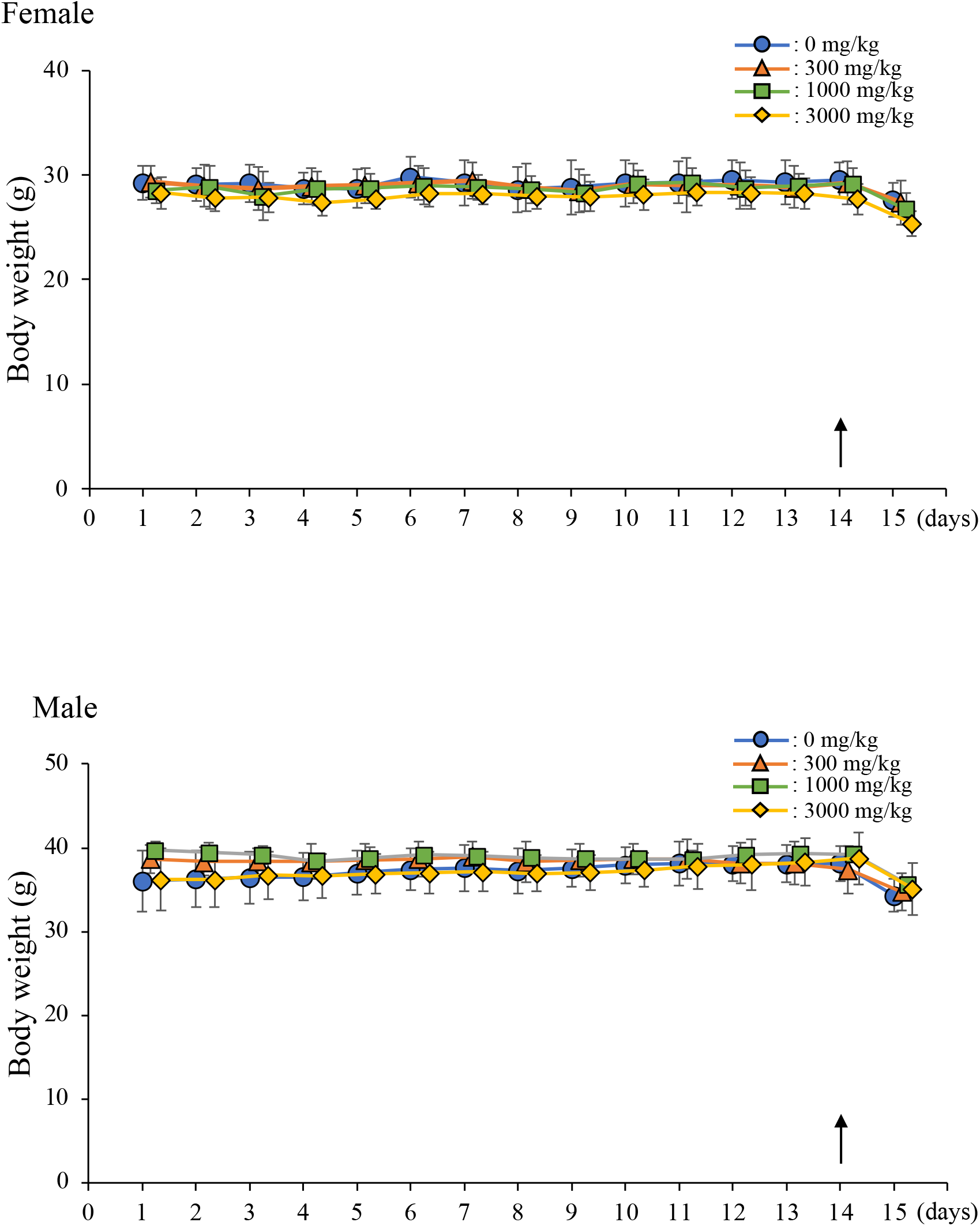
Body weight changes in male and female mice in the 14-day oral dose toxicity study. Values represent mean ± SD. Arrow indicates start of fasting. Females: n = 5 for all groups. Males: control group, n = 8; 300 mg/kg group, n = 5; 1,000 mg/kg group, n = 4; and 3,000 mg/kg group, n = 9.

In the 90-day oral toxicity study, there was no difference in body weight change during the 90-day treatment period in the 300, 1,000, and 3,000 mg/kg cricket powder groups compared with the control group for both males and females (Fig. 2). Furthermore, there were no differences in blood biochemistry (Table 6), blood parameters (Table 7), or organ weights as a percentage of total body weight (Table 8) in the 300, 1000, and 3,000 mg/kg cricket powder groups compared with the control group. These results suggest that there is no acute or chronic toxicity to individual mice following 14 and 90-day consumption of House cricket powder (≤3,000 mg/kg).

**Table 6.** Serum biochemical values of mice treated with House cricket powder for 90 days.

| Female |  |  |  |  |
| --- | --- | --- | --- | --- |
|  | Group |  |  |  |
|  | Control | 300 mg/kg | 1000 mg/kg | 3000 mg/kg |
| ALT<sup>1</sup> (IU/L) | 44 ± 14 | 40 ± 13 | 35 ± 9 | 38 ± 15 |
| BUN<sup>2</sup> (mg/dL) | 25.9 ± 4.6 | 21.8 ± 5.9 | 19.5 ± 3.5 | 24.6 ± 7.7 |
| CRE<sup>3</sup> (mg/dL) | 0.47 ± 0.28 | 0.87 ± 0.39 | 0.71 ± 0.42 | 0.74 ± 0.36 |
| CHO<sup>4</sup> (mg/dL) | 73 ± 29 | 62 ± 7 | 64 ± 9 | 73 ± 14 |
| TG<sup>5</sup> (mg/dL) | 72 ± 42 | 47 ± 20 | 56 ± 20 | 48 ± 23 |
| ALB<sup>6</sup> (g/dL) | 4.5 ± 0.9 | 4.7 ± 0.8 | 4.8 ± 0.5 | 4.2 ± 0.5 |
| CPK<sup>7</sup> (U/L) | 124 ± 61 | 128 ± 81 | 121 ± 52 | 99 ± 63 |
| Ca<sup>8</sup> (mg/dL) | 6.1 ± 2.2 | 4.9 ± 0.5 | 5.0 ± 0.5 | 5.3 ± 0.7 |
| Mg<sup>9</sup> (mg/dL) | 3.4 ± 0.6 | 2.9 ± 0.4 | 2.9 ± 0.5 | 3.0 ± 0.4 |
| Cl<sup>10</sup> (mmol/L) | 101 ± 4 | 99 ± 5 | 96 ± 4 | 100 ± 4 |

| Male |  |  |  |  |
| --- | --- | --- | --- | --- |
|  | Group |  |  |  |
|  | Control | 300 mg/kg | 1000 mg/kg | 3000 mg/kg |
| ALT<sup>1</sup> (IU/L) | 35 ± 11 | 37 ± 19 | 32 ± 7 | 33 ± 8 |
| BUN<sup>2</sup> (mg/dL) | 24.4 ± 4.1 | 27.5 ± 9.3 | 22.8 ± 4.0 | 24.1 ± 6.8 |
| CRE<sup>3</sup> (mg/dL) | 0.51 ± 0.22 | 0.66 ± 0.36 | 0.74 ± 0.20 | 0.77 ± 0.22 |
| CHO<sup>4</sup> (mg/dL) | 90 ± 12 | 87 ± 17 | 83 ± 17 | 77 ± 13 |
| TG<sup>5</sup> (mg/dL) | 83 ± 32 | 79 ± 25 | 73 ± 27 | 59 ± 20 |
| ALB<sup>6</sup> (g/dL) | 3.5 ± 1.0 | 3.7 ± 0.9 | 4.5 ± 0.5 | 4.4 ± 0.7 |
| CPK<sup>7</sup> (U/L) | 106 ± 28 | 178 ± 146 | 108 ± 63 | 87 ± 46 |
| Ca<sup>8</sup> (mg/dL) | 7.1 ± 1.8 | 7.1 ± 2.2 | 5.7 ± 1.2 | 5.4 ± 0.8 |
| Mg<sup>9</sup> (mg/dL) | 3.6 ± 0.7 | 3.7 ± 0.8 | 3.2 ± 0.8 | 3.0 ± 0.4 |
| Cl<sup>10</sup> (mmol/L) | 104 ± 5 | 104 ± 3 | 102 ± 3 | 100 ± 4 |

Values represent mean ± SD.
1: Aspartate aminotransferase, 2: Blood urea nitrogen, 3: creatinine, 4: total cholesterol, 5: triglycerides, 6: albumin, 7: creatine phosphokinase, 8: calcium, 9: magnesium, and 10: chloride.
Females: control, 300, and 3,000 mg/kg groups, n = 10; 1,000 mg/kg group, n = 9. Males: Control and 300 mg/kg groups, n = 11; 1,000 and 3,000 mg/kg groups, n = 10.

**Table 7.** Hematological values of mice treated with House cricket powder for 90 days.

| Female |  |  |  |  |
| --- | --- | --- | --- | --- |
|  | Group |  |  |  |
|  | Control | 300 mg/kg | 1000 mg/kg | 3000 mg/kg |
| WBC<sup>1</sup> (10<sup>3</sup>/μL) | 3.78 ± 1.99 | 4.48 ± 2.86 | 3.29 ± 1.65 | 5.91 ± 4.22 |
| RBC<sup>2</sup> (10<sup>6</sup>/μL) | 8.28 ± 0.81 | 8.53 ± 0.68 | 8.15 ± 0.89 | 8.19 ± 0.67 |
| Hemoglobin (g/dL) | 12.66 ± 1.31 | 13.01 ± 0.95 | 12.51 ± 1.15 | 12.60 ± 1.19 |
| Hematocrit (%) | 39.95 ± 4.23 | 41.78 ± 3.70 | 40.11 ± 3.90 | 39.76 ± 3.29 |
| MCV<sup>3</sup> (μm<sup>3</sup>) | 48.25 ± 1.37 | 48.93 ± 1.68 | 49.24 ± 1.30 | 48.57 ± 1.89 |
| MCH<sup>4</sup> (pg) | 15.30 ± 0.46 | 15.25 ± 0.55 | 15.39 ± 0.33 | 15.41 ± 0.73 |
| MCHC<sup>5</sup> (g/dL) | 31.73 ± 0.35 | 31.18 ± 0.66 | 31.24 ± 0.57 | 31.69 ± 0.60 |
| RDW<sup>6</sup> (%) | 13.66 ± 0.65 | 13.64 ± 0.63 | 13.57 ± 0.44 | 13.76 ± 0.51 |
| Platelets (10<sup>3</sup>/μL) | 69.5 ± 42.0 | 170.1 ± 109.0 | 74.4 ± 25.5 | 221.1 ± 195.5 |
| Lymphocyte (10<sup>3</sup>/μL) | 1.58 ± 0.79 | 1.60 ± 0.78 | 1.23 ± 0.62 | 2.44 ± 1.48 |
| Monocyte (10<sup>3</sup>/μL) | 0.09 ± 0.06 | 0.11 ± 0.08 | 0.07 ± 0.05 | 0.14 ± 0.13 |
| Granulocyte (10<sup>3</sup>/μL) | 2.11 ± 1.60 | 2.76 ± 2.01 | 1.99 ± 1.04 | 3.33 ± 2.73 |
| Eosinocyte (10<sup>3</sup>/μL) | 1.04 ± 0.91 | 1.39 ± 1.07 | 0.97 ± 0.61 | 1.33 ± 1.04 |

| Male |  |  |  |  |
| --- | --- | --- | --- | --- |
|  | Group |  |  |  |
|  | Control | 300 mg/kg | 1000 mg/kg | 3000 mg/kg |
| WBC<sup>1</sup> (10<sup>3</sup>/μL) | 6.13 ± 2.31 | 5.68 ± 2.72 | 4.90 ± 3.01 | 6.83 ± 3.97 |
| RBC<sup>2</sup> (10<sup>6</sup>/μL) | 8.23 ± 1.94 | 8.44 ± 0.52 | 8.97 ± 0.76 | 8.64 ± 0.43 |
| Hemoglobin (g/dL) | 12.54 ± 3.02 | 12.95 ± 0.32 | 13.36 ± 1.12 | 13.07 ± 0.65 |
| Hematocrit (%) | 39.94 ± 9.47 | 41.33 ± 1.83 | 43.21 ± 3.95 | 41.53 ± 2.11 |
| MCV<sup>3</sup> (μm<sup>3</sup>) | 48.45 ± 1.49 | 49.04 ± 1.73 | 48.18 ± 1.77 | 48.07 ± 1.09 |
| MCH<sup>4</sup> (pg) | 15.19 ± 0.47 | 15.39 ± 0.59 | 14.93 ± 0.43 | 15.10 ± 0.35 |
| MCHC<sup>5</sup> (g/dL) | 31.35 ± 0.36 | 31.38 ± 0.81 | 30.99 ± 0.46 | 31.43 ± 0.41 |
| RDW<sup>6</sup> (%) | 13.47 ± 0.23 | 13.75 ± 0.43 | 13.98 ± 0.75 | 13.84 ± 0.53 |
| Platelets (10<sup>3</sup>/μL) | 72.6 ± 31.9 | 195.1 ± 146.8 | 165.5 ± 108.9 | 172.9 ± 74.8 |
| Lymphocyte (10<sup>3</sup>/μL) | 2.34 ± 0.96 | 2.19 ± 0.99 | 1.98 ± 0.78 | 2.54 ± 1.34 |
| Monocyte (10<sup>3</sup>/μL) | 0.17 ± 0.06 | 0.15 ± 0.09 | 0.09 ± 0.08 | 0.14 ± 0.10 |
| Granulocyte (10<sup>3</sup>/μL) | 3.62 ± 1.50 | 3.34 ± 2.02 | 2.84 ± 2.20 | 4.14 ± 2.56 |
| Eosinocyte (10<sup>3</sup>/μL) | 1.65 ± 0.76 | 1.43 ± 1.10 | 1.33 ± 1.09 | 1.99 ± 1.23 |

Values represent mean ± SD.
1: White blood cell, 2: RBC, 3: mean corpuscular volume, 4: mean corpuscular hemoglobin, 5: mean corpuscular hemoglobin concentration, and 6: RBC distribution width.
Females: control and 300 mg/kg groups, n = 9; 1,000 and 3,000 mg/kg groups, n = 7. Males: control group, n = 11, 300, 1,000, and 3,000 mg/kg groups, n = 8.

**Table 8.** Relative organ weight of mice treated with House cricket powder for 90 days.

| Female |  |  |  |  |
| --- | --- | --- | --- | --- |
| OW/BW (%) | Group |  |  |  |
|  | Control | 300 mg/kg | 1000 mg/kg | 3000 mg/kg |
| Brain | 1.6022 ± 0.1735 | 1.6414 ± 0.1326 | 1.6377 ± 0.0935 | 1.5923 ± 0.1642 |
| Pituitary gland | 0.0101 ± 0.0046 | 0.0079 ± 0.0037 | 0.0113 ± 0.0028 | 0.0102 ± 0.0034 |
| Submandibular gland | 0.5154 ± 0.0903 | 0.5234 ± 0.1173 | 0.5625 ± 0.1004 | 0.5635 ± 0.0482 |
| Thymus | 0.1514 ± 0.0515 | 0.1637 ± 0.0465 | 0.1506 ± 0.0310 | 0.1386 ± 0.0491 |
| Heart | 0.4782 ± 0.0339 | 0.5683 ± 0.2267 | 0.4736 ± 0.0396 | 0.4872 ± 0.0468 |
| Lung | 0.5884 ± 0.1012 | 0.7293 ± 0.2419 | 0.6528 ± 0.0718 | 0.5974 ± 0.1049 |
| Liver | 4.3728 ± 0.3460 | 4.1870 ± 0.3891 | 4.4669 ± 0.2441 | 4.4709 ± 0.2109 |
| Spleen | 0.3267 ± 0.0822 | 0.2943 ± 0.0859 | 0.2648 ± 0.0353 | 0.2893 ± 0.0329 |
| Pancreas | 0.6797 ± 0.1466 | 0.6700 ± 0.1521 | 0.6249 ± 0.2381 | 0.6598 ± 0.0824 |
| Adrenal gland (R) | 0.0159 ± 0.0035 | 0.0147 ± 0.0038 | 0.0146 ± 0.0031 | 0.0146 ± 0.0029 |
| Kidney (R) | 0.6733 ± 0.0843 | 0.6967 ± 0.0729 | 0.7045 ± 0.0420 | 0.6751 ± 0.0693 |
| Adrenal gland (L) | 0.0186 ± 0.0046 | 0.0156 ± 0.0028 | 0.0173 ± 0.0056 | 0.0150 ± 0.0042 |
| Kidney (L) | 0.6620 ± 0.0741 | 0.6494 ± 0.0464 | 0.6676 ± 0.0370 | 0.6579 ± 0.0343 |
| Ovary (R) | 0.0269 ± 0.0121 | 0.0387 ± 0.0141 | 0.0294 ± 0.0081 | 0.0324 ± 0.0138 |
| Ovary (L) | 0.0269 ± 0.0102 | 0.0375 ± 0.0119 | 0.0284 ± 0.0110 | 0.0349 ± 0.0130 |
| Uterus | 0.5445 ± 0.2399 | 0.7079 ± 0.3909 | 0.6093 ± 0.1383 | 0.5654 ± 0.2798 |
| Visceral fat (R) | 0.3507 ± 0.3456 | 0.1484 ± 0.1894 | 0.4934 ± 0.2432 | 0.2581 ± 0.2209 |
| Visceral fat (L) | 0.3005 ± 0.2602 | 0.1341 ± 0.1394 | 0.4212 ± 0.2342 | 0.2226 ± 0.1945 |
| Subcutaneous fat | 0.4042 ± 0.1881 | 0.4060 ± 0.1888 | 0.4911 ± 0.2221 | 0.3361 ± 0.1402 |
| Gastrocnemius muscle (R) | 0.5669 ± 0.1381 | 0.5793 ± 0.1788 | 0.4767 ± 0.0718 | 0.5529 ± 0.1666 |
| Gastrocnemius muscle (L) | 0.5645 ± 0.1223 | 0.5040 ± 0.1073 | 0.4865 ± 0.0642 | 0.5733 ± 0.1061 |
| Soleus muscle (R) | 0.0308 ± 0.0150 | 0.0277 ± 0.0153 | 0.0319 ± 0.0116 | 0.0362 ± 0.0194 |
| Soleus muscle (L) | 0.0300 ± 0.0116 | 0.0322 ± 0.0157 | 0.0292 ± 0.0105 | 0.0299 ± 0.0107 |

Male
| OW/BW (%) | Group |  |  |  |
| --- | --- | --- | --- | --- |
|  | Control | 300 mg/kg | 1000 mg/kg | 3000 mg/kg |
| Brain | 1.3297 ± 0.1145 | 1.3096 ± 0.1000 | 1.2957 ± 0.1230 | 1.3680 ± 0.0878 |
| Pituitary gland | 0.0074 ± 0.0032 | 0.0068 ± 0.0022 | 0.0059 ± 0.0023 | 0.0061 ± 0.0027 |
| Submandibular gland | 0.7523 ± 0.1030 | 0.7338 ± 0.0868 | 0.7665 ± 0.0566 | 0.6927 ± 0.1325 |
| Thymus | 0.0915 ± 0.0238 | 0.0965 ± 0.0289 | 0.0912 ± 0.0168 | 0.0948 ± 0.0372 |
| Heart | 0.5079 ± 0.0551 | 0.5067 ± 0.0560 | 0.5127 ± 0.0667 | 0.5176 ± 0.0433 |
| Lung | 0.5324 ± 0.0428 | 0.6463 ± 0.1577 | 0.6136 ± 0.0963 | 0.6110 ± 0.1748 |
| Liver | 4.3486 ± 0.1905 | 4.2449 ± 0.1357 | 4.3052 ± 0.1660 | 4.2010 ± 0.3310 |
| Spleen | 0.2115 ± 0.0426 | 0.2421 ± 0.0512 | 0.2444 ± 0.0378 | 0.2030 ± 0.0357 |
| Pancreas | 0.6455 ± 0.1626 | 0.5703 ± 0.2285 | 0.6348 ± 0.1322 | 0.5597 ± 0.2207 |
| Adrenal gland (R) | 0.0080 ± 0.0035 | 0.0108 ± 0.0067 | 0.0087 ± 0.0030 | 0.0092 ± 0.0022 |
| Kidney (R) | 0.8807 ± 0.0724 | 0.9136 ± 0.0690 | 0.8607 ± 0.1123 | 0.8933 ± 0.1368 |
| Adrenal gland (L) | 0.0097 ± 0.0040 | 0.0103 ± 0.0038 | 0.0096 ± 0.0030 | 0.0099 ± 0.0025 |
| Kidney (L) | 0.8264 ± 0.0784 | 0.8369 ± 0.0871 | 0.8130 ± 0.0921 | 0.8502 ± 0.1138 |
| Testis (R) | 0.4466 ± 0.0595 | 0.4663 ± 0.0681 | 0.4660 ± 0.1003 | 0.4642 ± 0.0609 |
| Testis (L) | 0.4335 ± 0.0593 | 0.4474 ± 0.0658 | 0.4801 ± 0.1513 | 0.4312 ± 0.0669 |
| Seminal vesicle/Coagulating gland | 1.0256 ± 0.1143 | 0.9737 ± 0.1812 | 0.8970 ± 0.1625 | 0.9878 ± 0.1792 |
| Visceral fat (R) | 0.4935 ± 0.2093 | 0.4655 ± 0.1832 | 0.3933 ± 0.1142 | 0.3776 ± 0.1858 |
| Visceral fat (L) | 0.5074 ± 0.2108 | 0.4579 ± 0.1760 | 0.3720 ± 0.1363 | 0.4263 ± 0.2491 |
| Subcutaneous fat | 0.4686 ± 0.1671 | 0.4224 ± 0.2143 | 0.4585 ± 0.1960 | 0.4046 ± 0.1345 |
| Gastrocnemius muscle (R) | 0.5347 ± 0.1792 | 0.4786 ± 0.1687 | 0.5424 ± 0.0954 | 0.5091 ± 0.0999 |
| Gastrocnemius muscle (L) | 0.5173 ± 0.0840 | 0.5163 ± 0.1147 | 0.5051 ± 0.1234 | 0.4697 ± 0.0649 |
| Soleus muscle (R) | 0.0417 ± 0.0226 | 0.0474 ± 0.0213 | 0.0460 ± 0.0259 | 0.0870 ± 0.1450 |
| Soleus muscle (L) | 0.0386 ± 0.0174 | 0.0472 ± 0.0235 | 0.0362 ± 0.0207 | 0.0772 ± 0.1127 |

Values represent mean ± SD.
OW, organ weight, BW, body weight.

**Figure 2.**
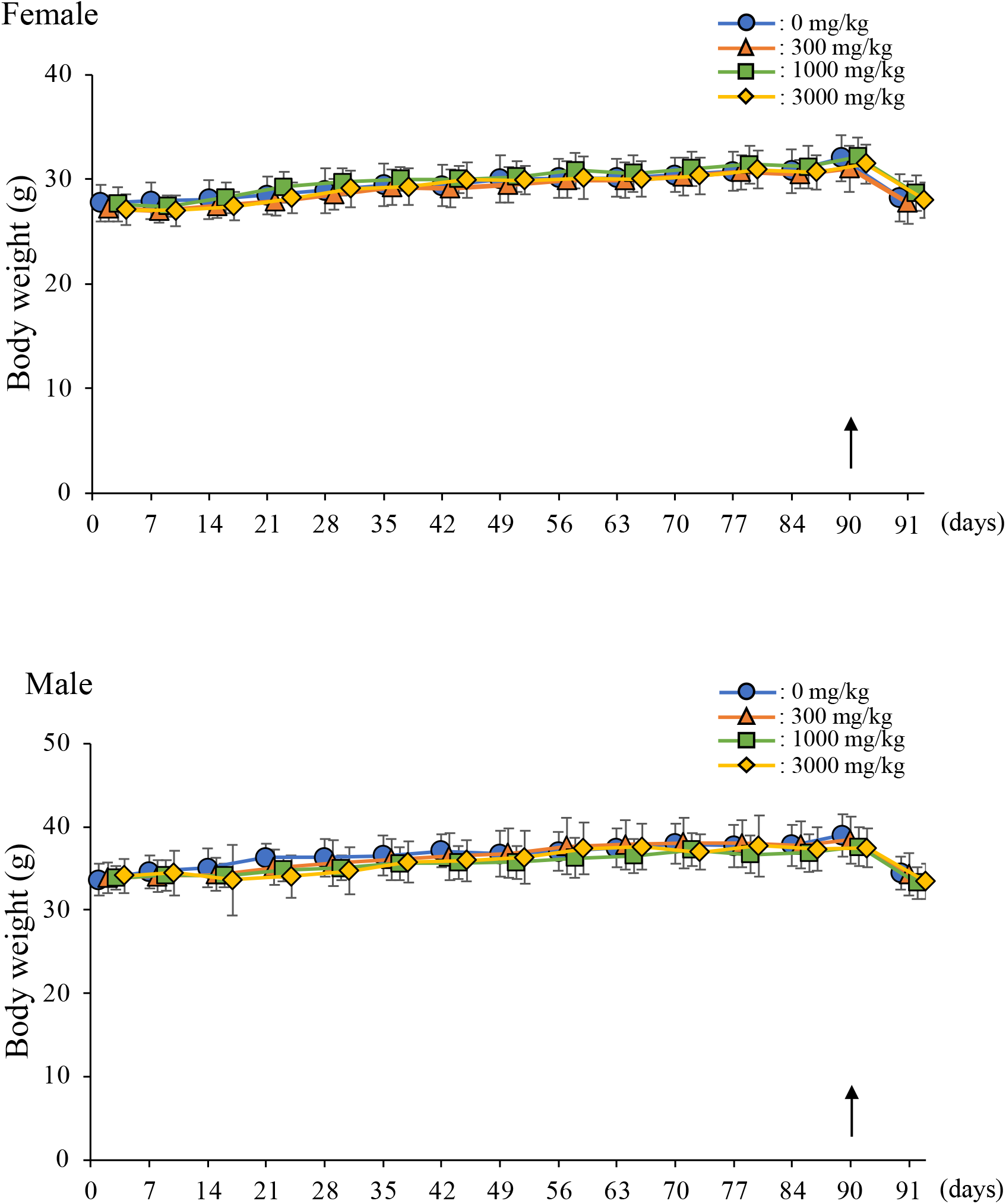
Body weight changes in male and female mice in the 90-day oral dose toxicity study. Values represent mean ± SD. Arrow indicates start of fasting. Females: control and 300 mg/kg groups, n = 12; 1,000 mg/kg group, n = 9; 3,000 mg/kg group, n = 10. Males: control, 300 mg/kg, and 3,000 mg/kg groups, n = 11; 1,000 mg/kg group, n = 10.

## Discussion

Insects are used as food in many parts of the world, including Asia, North and South America, and Africa. In Japan, bee larvae and locusts are also used as food and it is estimated that 1,400 species of insects are used as food worldwide. Other insects, such as yellow mealworm (*Tenebrio molitor* larva) and *Gryllus bimaculatus*, have been evaluated for their safety as food (Han *et al*., 2014; Turck, *et al*., 2021b; Ryu *et al*., 2016). Although house crickets (*Acheta domesticus*) have been reported to be safe as food and their risk has been evaluated (Turck, *et al*., 2021a; Fernandez-Cassi *et al*., 2019), their cytotoxicity and toxicity to organisms have not been evaluated in accordance with the OECD guidelines. The present study evaluated the cellular genotoxicity of cricket powder using an *in vitro* chromosome aberration test and its genetic, acute, and chronic biological toxicity in ICR mice in accordance with the OECD guidelines.

*In vitro* chromosomal aberration test showed no clear cytotoxicity under treatment of 5,000 µg/mL cricket powder on CHL-IU cells (Table 1). *In vivo* micronucleus test results showed no genotoxic effects of ingestion of ≤2,000 mg/kg cricket powder on erythrocytes in mouse bone marrow (Table 2). However, the present study was unable to determine the genotoxicity of >2,000 mg/kg cricket powder due to ingestion; therefore, further studies are required to evaluate the toxicity of ingestion of higher concentrations of cricket powder. In the repeated oral toxicity study, no significant changes in body weight were observed in males and females following 14 and 90-day continuous administration studies of cricket powder (≤3,000 mg/kg) (Figs. 1, 2). Furthermore, 14-day administration of cricket powder did not result in weight gain over time in females and males in any of the groups. This lack of weight gain may result from the stress of repeated forced oral administration. In addition, no abnormalities were observed in blood properties, blood biochemistry tests, or organ weights in both males and females after administration of cricket powder (≤3,000 mg/kg) for 14 and 90 days (Tables 3-8). In conclusion, the present study found no cellular or biological toxicity from consumption of House crickets at concentrations ≤3,000 mg/kg, indicating that House crickets may be a safe and high-quality animal protein foodstuff. This study is useful evidence for evaluating the toxicity of House crickets *in vivo*. Future studies are planned to re-evaluate AST, ALP, T-BIL, UA, GLU, TP, LDH, IP, Na, and K, which could not be measured in the blood biochemistry analysis in the 90-day oral toxicity study since the blood samples were hemolyzed. In addition, there is concern that adult crickets, such as the House cricket and Desert locust, contain substantial amounts of purines, which may aggravate diseases, such as gout (Sabolová *et al*., 2021); therefore, we will conduct component analysis and safety evaluation using House crickets before sexual maturity to determine the most suitable crickets to harvest for use as foodstuffs.

## Acknowledgments

This work was supported by Cabinet Office, Government of Japan, Cross-ministerial Moonshot Agriculture, Forestry and Fisheries Research and Development Program, “Technologies for Smart Bio-industry and Agriculture” (JPJ009237).

## Conflict of interest

There are no conflicts of interest to declare.

## Notes

### Competing Interest Statement

The authors have declared no competing interest.

